# IL-27 Reduces Cell-Surface LDLR in Human Macrophages and Restricts VSV-G-Pseudotyped HIV infection

**DOI:** 10.64898/2026.10.22.761012

**Authors:** Qian Chen, Udeshika Kariyawasam, Mayra Maruquez, Jordan Metz, Jun Yang, Adam Rupert, Tomozumi Imamichi

## Abstract

The low-density lipoprotein receptor (LDLR) is a major cellular receptor for vesicular stomatitis virus glycoprotein (VSV-G), which is widely used to pseudotyped virus vectors. A replication incompetent VSV-G-pseudotyped HIV (VSV-G-HIVLuc) has been used in HIV infection studies. Although IL-27-polarized macrophages induce multiple antiviral genes and resist replication-competent HIV infection, their impact on VSV-G-HIVLuc infection has not been studied. Here, we evaluated the role of IL-27 polarization on VSV-G-HIVLuc infection. Infection was monitored by luciferase activity. Cell-surface LDLR was analyzed by flow cytometry. LDLR transcripts and protein expression were quantified by qRT-PCR and Western blotting, respectively. Viral binding was assessed by measuring cell-associated p24 antigen, while post-entry events were evaluated by quantifying late reverse-transcription products. VSV-G-HIVLuc infection in IL-27-polarized M-CSF-induced macrophages (27M-Mac) and IL-27-polarized GM-CSF-induced macrophages (27GM-Mac) was reduced to 5.8±0.4% and 5.9±2.1%, respectively (both *p*<0.0001), compared to unpolarized macrophages. 27M-Mac and 27GM-Mac reduced cell-surface LDLR to 38.3±2.6% (*p*<0.01) and 31.4±3.2% (*p*<0.01). Western blotting showed that total LDLR protein decreased to 48±0.8% in 27M-Mac (*p*<0.01), whereas 27GM-Mac showed no significant change. However, LDLR transcripts increased approximately 1.6-1.9-fold following IL-27 treatment in both macrophage types (*p*<0.01). Viral binding to 27M-Mac decreased modestly, while binding to 27GM-Mac decreased significantly to 67.5 ± 7.5% of that to GM-Mac (*p*<0.05, n=3). In contrast, late reverse-transcription products decreased to 23±2.1% (p<0.0001) and 44±1.5% (p<0.0001). These findings demonstrate that IL-27 downregulates cell-surface LDLR expression in macrophages despite increasing LDLR transcript abundance, suggesting that IL-27 restricts VSV-G-pseudotyped Lentiviral infection through both receptor-associated and additional post-entry mechanisms.

## Introduction

Interleukin-27 (IL-27) is a heterodimeric cytokine composed of the p28 (also known as IL-30) and Epstein-Barr virus-induced gene 3 subunits [1]. IL-27 signals through IL27 receptor composed of WSX-1 and gp130 and activates predominantly STAT1 and STAT3 signaling pathways. Accordingly, IL-27 is classified as a member of the IL-6/gp130 cytokine family, and regulates diverse cell types, including monocytes, T cells, macrophages, dendritic cells, hepatocytes, keratinocytes, and endothelial cells [2-5].

Macrophages display substantial phenotypic and functional plasticity in response to environmental signals. Human monocytes-differentiated with GM-CSF or M-CSF are commonly used as M1-like (GM-Mac) and M2-like (M-Mac) macrophage models, respectively [6-9]. We previously demonstrated that IL-27 further polarizes both macrophage populations, generating IL-27-treated GM-Mac (27GM-Mac) and M-Mac (27M-Mac) with convergent phenotypic and functional characteristics [9,10]. Single-cell RNA sequencing revealed substantial convergence of their gene-expression profiles, and both IL-27-polarized macrophage populations exhibited enhanced antiviral activity.

Recently, IL-6 was reported to downregulate low-density lipoprotein receptor (LDLR) expression in endothelial cells [11], raising the possibility that regulation of LDLR may extend to other members of the IL-6/gp130 cytokine family. LDLR serves as a major cellular receptor for vesicular stomatitis virus glycoprotein (VSV-G) [12,13]. Because of its broad cellular tropism, VSV-G is widely used to pseudotyped lentiviral and retroviral vectors for gene delivery [14,15]. We previously used a replication-incompetent VSV-G-pseudotyped HIV vector encoding luciferase (VSV-G-HIVLuc) and observed markedly reduced infection in IL-27-diffrentiated macrophages [16]. However, the mechanism underlying this reduced susceptibility to VSV-G-HIVLuc remained unknown. In the present study, we investigated whether IL-27 regulates LDLR expression in human macrophages and whether such regulation contributes to the reduced susceptibility of IL-27-polarized macrophages to VSV-G-pseudotyped HIV. We examined cell-surface LDLR expression, and total LDLR protein, LDLR transcription, VSV-G HIVLuc binding, and reverse transcript copy number and virus production in M-, GM-, 27M-, and 27GM-Mac. Our findings reveal that IL-27 selectively reduces cell-surface LDLR with decreasing total LDLR expression in M-Mac but not in GM-Mac and provide a potential mechanism linking IL-27-mediated macrophage polarization to reduced VSV-G-dependent transduction.

## Materials and Methods

### Ethics statement

All experimental procedures were approved by the National Cancer Institute at Frederick (the protocol code number: 2016–19A6/11-30) and performed in accordance with the relevant guidelines and regulations.

### Cells, Viruses and Reagents

CD14+ monocytes were isolated from health donors’ PBMCs and differentiated into M-Mac and GM-Mac using M-CSF (R&D Systems Minneapolis, MN, USA) and GM-CSF (R&D Systems), respectively, as previously described [10]. M-Mac and GM-Mac cells were polarized to 27M-Mac and 27GM-Mac by culturing for 3-days in the presence of 100 ng/mL IL-27 (R&D Systems) in completed D-MEM medium (Thermo Fisher Scientific, Waltham, MA, USA) supplemented with 10%-FBS (R&D Systems), gentamycin (Thermo Fisher) and HEPES (Quality Biology, Gaithersburg, MD) [10]. VSVG-HIVLuc virus was produced by co-transfecting HEK293T(ATCC, Manassas, VA, USA) with pHIVNL4-3ΔEnv/Vpr Luciferase and pVSV-G plasmids and concentrated using ultracentrifugation as describe previously [16]. Virus p24 concentration was determined by quantification of p24 antigen using Alliance HIV-1 p24 Antigen ELISA Kit (Revvity, Waltham, MA, USA) [9].

### Detection of LDLR on cell surface

Cell-surface LDLR expression was quantified by flow cytometry as previously described [9]. Briefly, cells were pretreated with Fcγ Receptor Blocker (Innovex Biosciences, Richmond, CA, USA) and subsequently incubated with either an APC-conjugated mouse IgG1 isotype control (R&D Systems, Cat# IC002A) or APC-conjugated anti-human LDLR antibody (R&D Systems, Cat# FAB2148A). Stained cells were analyzed using an LSR Fortessa flow cytometer (BD Biosciences, San Jose, CA, USA). Flow cytometry data were analyzed using FlowJo software version 10.10.1 (BD Biosciences).

### Western Blotting

Western blotting (WB) was performed using total cell lysate in RIPA lysis buffer as previously described [10]. Mouse anti-LDLR (Abcam # ab271189) or mouse anti-β-Actin (Millipore Sigma, Cat #A5316) were used to detect intended proteins. Bands were detected with HRP-conjugated Goat anti-mouse Ig with ECL Prime Western Blotting Detection Reagent (Cytiva Life Sci-ences, Marlborough, MA, USA) and imaged with an Azure 300 system (Azure Biosystems, Dublin, CA, USA). Band intensities were quantified using Fiji Version 2.16 (National Institutes of Health, Bethesda, MD, USA) [10].

### Quantitative RT-PCR (qRT-PCR)

Total cellular RNA was isolated using the RNeasy Kit (Qiagen), and cDNA was synthesized using TaqMan reverse transcription reagents (Thermo Fisher Scientific) as described previously [10]. mRNA levels of genes of interest were measured by qRT-PCR on a CFX96 real-time system (Bio-Rad, Hercules, CA, USA) as described previously [10]. GAPDH was used as a reference for normalization. All probes were purchased from Thermo Fisher Scientific.

### Single-cell RNA-Seq analysis

Single-cell RNA libraries were constructed using the Chromium Next GEM Single Cell 5’ Reagent Kit V2 (Dual Index) (10x Genomics, Pleasanton, CA, USA) as described before [10] and libraries were run in NextSeq 2000. Sequence data were processed using Cell Ranger v6.0.1 (10x Genomics) [10]. Genes were considered differentially expressed at *p* < 0.05 with an absolute fold change ≥3. To adjust *p* values, Tukey’s Honestly Significant Difference (HSD) is used to compare all possible pairs of group means. The statistical significance threshold was set as p-value <0.05 and |fold change| ≥3. Genes exceeding these combined criteria were color-coded, significantly up-regulated in red and significantly down-regulated genes in blue, while non-significant genes remained black. Genes of interest were labeled with its gene name.

### VSVG-HIVLuc binding assay

M-, GM-, 27M- or 27GM-Mac (1×10^6^ cells) were incubated with 100 ng p24/ml of VSVG-HIVLuc in 1.5 mL tube on a rotator for 2 h at 4°C. Cells were washed with cold PBS without vortexing and then were lysed using 100 µL of Radioimmunoprecipitation assay (RIPA) buffer (Boston BioProducts, Milford, MA, USA) supplemented with proteinase and phosphatase inhibitors cocktail (Thermo Fisher Scientific) at 4 C for 15 min. Cell lysate was collected and cell debris free lysate was prepared by 3500 rpm for 10 min at 4°C. Bound virus on cells were assessed by quantification of p24 amounts using p24 antigen capture assay kit and for normalization, total cellular protein was measured using BCA protein assay kit (Thermo Fisher Scientific).

### VSVG-HIVLuc infection assay

VSVG-HIVLuc infection assay was performed in 96-well plates as previously described. M-, GM-, 27M-, or 27GM-Mac (50x10^3^ cells /well) in 96-well plates were infected with 100 ng p24/mL of VSVG HIVLuc for 2 h at 37°C. Infected cells were cultured for 2 days. Infection was quantified with measuring luciferase activity using Bright Glo Luciferase assay system (Promega, Madison, WI, USA) as described in previously [16]. All replication assays were performed in quadruplicate. Luciferase actively was detected Spark microplate reader (Tecan, Morrisville, NC, USA).

### Copy number assay

Proviral HIV copy number was quantified using the QuantStudio Absolute Q Digital PCR (dPCR) System (Thermo Fisher Scientific). Host genomic DNA was extracted from VSVG-HIVLuc-infected M-, GM-, 27M or 27GM-Mac using QIAamp DNA Blood Kits (Qiagen, Germantown, MD, USA). Total 100 ng genomic DNAs were subject of the PCR assay. DNA was mixed with 2 µL of Absolute Q Universal DNA Digital PCR Master Mix (Thermo Fisher Scientific) and 0.5 µL of 20X dPCR probe (Thermo Fisher Scientific) in total 12 µL was loaded into the MAP plate (Thermo Fisher Scientific) and then ran using a default program for copy number assay. Assay was conducted triplicate and then data was normalized using copy number of RNAP gene. Custm made HIV gag probe was purchased from Thermo Fisher Scientific.

### Statistical Analysis

Intergroup comparisons were performed using one-way ANOVA (GraphPad, San Diego, CA, USA). p-values <0.05 were considered statistically significant: ^*^ p < 0.05, ^* *^ p < 0.01, ^* * *^ p < 0.001, ^* * * *^ p < 0.0001; p > 0.05 was considered not significant (ns).

## Results

### Comparison of LDLR expression among four macrophages

We compared cell-surface LDLR expression among M-, GM-, 27M-, and 27GM-Mac using flow cytometry. As shown in **Figures 1A and 1B**, surface LDLR expression in GM-Mac was comparable that in M-Mac. IL-27 polarization reduced surface LDLR expression in 27M-Mac and 27GM-Mac to 38.3 ± 2.6% (p<0.01) and 31.4 ± 3.2% (p<0.01) of that in M-Mac and GM-Mac, respectively. IL-6-mediated downregulation of LDLR has been reported to involve lysosomal degradation of LDLR protein [11]. Therefore, we next examined whether the reduction in cell-surface LDLR was associated with decrease in total cellular LDLR protein levels. Western blot analysis revealed that total LDLR protein levels decreased by approximately 50% in 27M-Mac compared with M-Mac (*p* < 0.01). Notably, total LDLR protein levels in GM-Mac were approximately 50% lower than those in M-Mac (*p* < 0.001), whereas no significant difference was observed between GM-Mac and 27GM-Mac **(Figures 1C and 1D)**. To further investigate whether these differences in total LDLR protein levels were associated with changes in LDLR mRNA expression, qRT-PCR analysis was performed. LDLR mRNA levels increased approximately 1.6-fold in 27M-Mac compared with M-Mac (*p* < 0.01). In 27GM-Mac, LDLR mRNA levels increased approximately 1.9-fold compared with M-Mac (*p* < 0.001) and 1.6-fold compared with GM-Mac (*p* < 0.001) **(Figure 1E)**.

**Figure 1.**
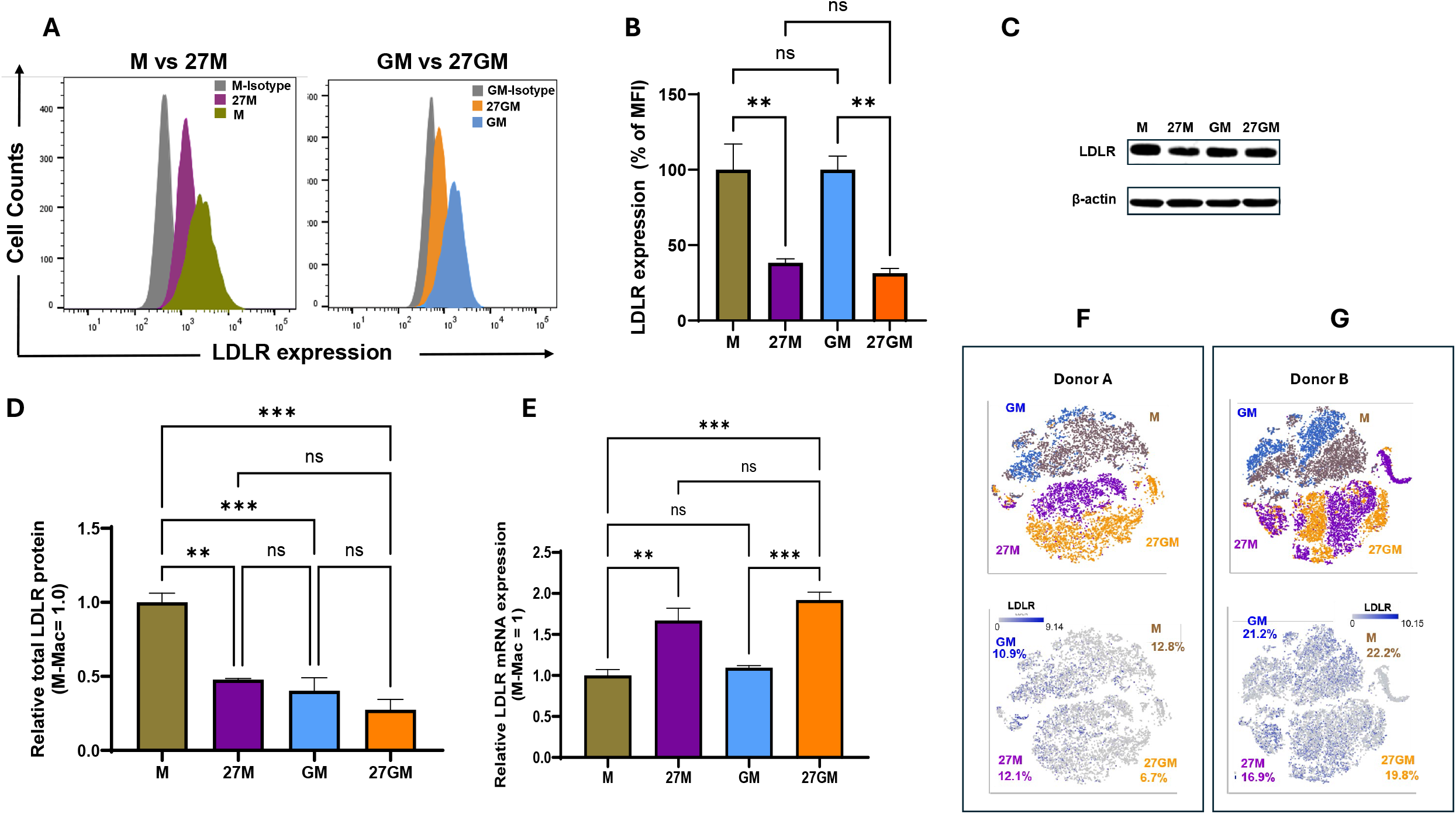
Characterization of IL-27 effect on LDLR expression. **(A and B)** Cell-surface LDLR expression was analyzed by flow cytometry. Each cell type was stained with an APC-conjugated isotype control or anti-human LDLR antibody and analyzed using an LSR Fortessa flow cytometer. Gray is isotype control. M-Mac (olive green), 27M-Mac (purple), GM-Mac (blue), and 27GM-Mac (orange). Representative data from one of four independent donors are shown (A). Flow cytometry data were analyzed using FlowJo software (version 10.10.1). Summary data are presented as means ± SE from four independent donors (B). **(C and D)** Total cellular lysates (20 μg of protein per sample) were subjected to Western blotting using an anti-LDLR antibody. After detecting LDLR band, antibody was stripped and then reprobed using anti β-actin antibody as the loading control. Representative results from one of three independent donors are shown (C). LDLR protein levels in M-, 27M-, GM- and 27GM-Mac were quantified by densitometric analysis using Fiji (version 2.16) from three independent donors. LDLR band intensities were normalized to the corresponding β-actin band intensities. Within each experiment, the normalized LDLR expression in M-Macs was set to 100%, and the expression in the other cell types was expressed relative to this value. Bars represent the mean ± SE of three independent experiments. (D). **(E)** LDLR mRNA levels in M-, 27M-, GM-, and 27GM-Mac were quantified by qRT-PCR using total cellular RNA, with *GAPDH* as the reference gene. Relative expression was calculated using the 2^−ΔΔCt method, with M-Mac as the calibrator (set to 1). Bars represent the mean ± SE of three independent experiments. Statistical comparisons were performed using ΔCt values. **(F and G)** Single-cell RNA sequencing was performed using freshly prepared M-, GM-, 27M-, and 27GM-Mac from two independent donors (donors A and B), as described in Materials and Methods. The upper t-SNE panels show the distribution of M-Mac (olive green), 27M-Mac (purple), GM-Mac (blue), and 27GM-Mac (orange). The lower panels show LDLR mRNA expression, with gray indicating low or undetectable expression and progressively darker blue indicating higher expression, as shown in the color scales. Percentages below each cell-type label indicate the proportion of LDLR mRNA-expressing cells within that population. Statistical analyses were conducted using ANOVA, ^*^ p < 0.05, ^* *^ p < 0.01, ^* * *^ p < 0.001, ns, not significant.

We have previously compared gene expression profile among these four macrophages subtypes using scRNA seq [9]. Here, we assessed distribution of LDLR-expressing cells in that dataset. As shown **Figure 1F and 1G**, the *t*-SNE plots showed that cells expressing LDLR mRNA were distributed across the macrophage populations, with no substantial change in their distribution following IL-27 polarization.

### IL-27-polarized macrophages exhibit VSVG binding, DNA accumulation and reporter expression

We previously reported that IL-27 polarizes M-Mac (M2-like macrophages) and GM-Mac (M1-like macrophages) toward convergent phenotypic states [9,10]. Having observed the downregulation of LDLR expression on cell surface, we next examined whether IL-27-mediated downregulation of cell-surface LDLR was associated with reduced viral binding. M-, GM-, 27M-, and 27GM-Mac were incubated with VSV-G-HIVLuc at 4°C, and cell-associated viral particles were quantified by HIV p24 ELISA.

Viral binding to GM-Mac was approximately 2.5-fold higher than that to M-Mac (*p* < 0.01) **(Figure 2A)**, despite comparable cell-surface LDLR expression between the two macrophage populations **(Figures 1A and 1B)**. Although viral binding to 27M-Mac decreased to 50% of that to M-Mac, this reduction was not statistically significant. In contrast, viral binding to 27GM-Mac decreased significantly to 67.5 ± 7.5% of that to GM-Mac (*p* < 0.05). These findings indicate that IL-27-mediated downregulation of LDLR expression is not consistently accompanied by significant reduction in VSVG-HIVLuc binding, suggesting that additional factors on cell surface may influence viral binding [12,13]. We next quantified HIV infection by measuring luciferase activity. Because luciferase gene is encoded within VSVG-HIVLuc construct [17] and its expression is driven by the HIV LTR activity, luciferase activity provides a functional measure of HIV LTR-driven gene expression. Luciferase activity in 27M- and 27GM-Mac was significantly reduced to nearly 6 % of that in the corresponding control cell type respectively (p<0.0001 for both cells) (**Figure 2B**). Because the magnitude of luciferase inhibition was substantially greater than the reduction in viral binding, we next quantified viral DNA copy number by measuring late reverse-transcription (RT) products using digital PCR (dPCR). Late RT products were reduced to 25% in 27M-Mac (p<0.0001) and 45 % in 27GM-Mac (p<0.0001) relative to their respective controls **(Figure 2C**). Those were not the same level of suppression in Luc activity **(Figure 2B)**, indicating that HIV production is further down regulated in 27M- and 27GM-Mac at transcription or translation step.

**Figure 2.**
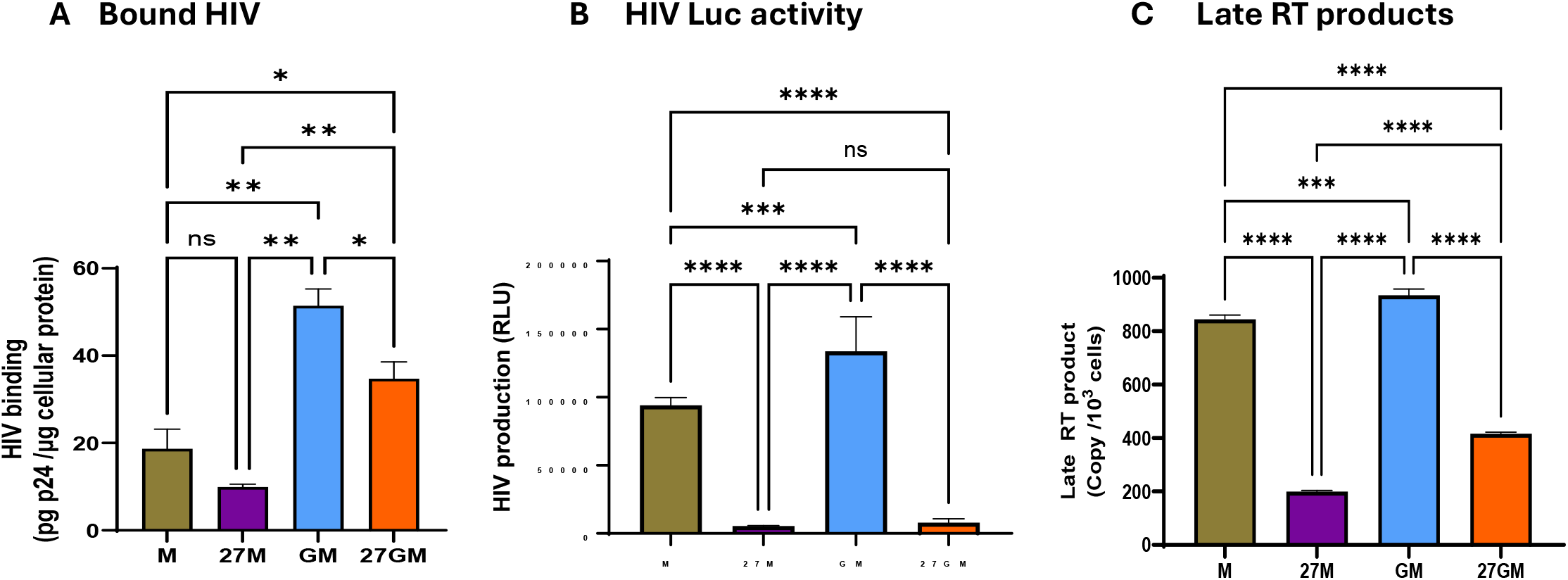
Viral binding and infection in macrophages with reduced cell-surface LDLR expression. **(A**) Quantification of VSV-G–HIVLuc binding. Each cell type was incubated with VSV-G– HIVLuc at 4°C and subsequently lysed in RIPA buffer. Cell-associated viral p24 antigen was quantified using an HIV-1 p24 antigen ELISA kit, and total cellular protein was measured using a BCA protein assay kit. The amount of bound virus was normalized to total cellular protein. Each assay was done quadruplicate. Representative data from one of three independent experiments. Data are presented as means ± SD. (**B**) HIV expression in each cell type was assessed by measuring luciferase activity. Each assay was done sextuplicate. Representative data from one of three independent experiments. Data are presented as means ± SD. (**C**) Later RT product was quantified as HIV DNA copy numbers using a dPCR assay as described in Materials and Methods. Representative data from one of three independent experiments. Each assay was done triplicate. Data are presented as means ± SD. Statistical analyses were conducted using ANOVA, ^*^ p < 0.05, ^* *^ p < 0.01, ns, not significant.

## Discussion

We have investigated IL-27-mediated macrophage polarization[9,10]. A recent study reported that IL-6 down regulates the expression of LDLR on endothelial cells [11]. As IL-27 is a member of IL-6/gp130 cytokine family, in the current study, we assessed the role of IL-27-mediated (a member of IL-6 family cytokine) polarization on the LDLR expression in M-1 and M2-like macrophages (those cells were designated GM- and M-Mac in the current study). As expected, IL-27 polarization down-regulated the expression of the receptor on both cell types. WB results demonstrated that total cellular LDLR amounts in M-Mac were higher than that in GM-Mac, indicating that M-Mac may have different mechanism in intracellular trafficking of LDLR [18], as reported in other receptors such as Fc gamma receptor [19], chemokine or cytokine receptors [20-22]. Although in our previous scRNA seq analysis, LDLR gene was not identified as a differentiated gene (as our cut off was ≥3-fold in absolute changes). interestingly, LDLR mRNA levels increased following IL-27 treatment, whereas total LDLR protein was significantly reduced in 27M-Mac but not in 27GM-Mac. This discrepancy suggests that IL-27 regulates LDLR expression beyond transcription. 27M-Mac may enhance protein degradation, similar to the mechanism previously reported in IL-6-treated cells [11].

IL-27 treatment reduced cell-surface LDLR expression in both macrophage populations; however, its effect on VSV-G–HIVLuc binding differed between the two populations. Viral binding was significantly reduced in 27GM-Mac but not in 27M-Mac, indicating that decreased surface LDLR expression alone does not necessarily correspond to a proportional reduction in viral binding to the cells. Furthermore, GM-Mac exhibited substantially higher viral binding than M-Mac despite comparable cell-surface LDLR expression. These observations suggest that receptor accessibility, membrane organization, or additional viral binding factors may influence VSV-G-mediated viral binding, since VSVG envelope proteins binds to other cellular proteins [13].

As shown previously, the IL-27 receptor (a complex of WSX1 and gp130) is not uniformly expressed across macrophages. FACS data indicated that less than 10% cells express both WSX1 and gp130 [10]. On contrary, in the current study, FACS data indicated that LDLR expressed nearly 85 % of M-Mac and GM-Mac. Thus, reduced surface LDLR expression reflects both direct responses to IL-27 and indirect effects mediated by factors released from IL-27-responsive cells. However, it is important to note that the percentages alone do not establish an indirect mechanism, since cells without detectable receptor staining may still respond to IL-27. The mechanism of the down regulation requires further investigation between IL27R expressing and non-expressing cells.

Luciferase activity was markedly suppressed in both 27M- and 27GM-Mac. Intracellular HIV DNA was also reduced, but to a lesser extent than luciferase activity. This difference suggests that additional restrictions affecting steps between HIV DNA accumulation and reporter expression may contribute to the antiviral phenotype. Potentially affected steps include nuclear import, integration, transcription, and translation; the present assays do not distinguish among them.

Taken together, these findings suggest that IL-27-polarized macrophages restrict VSV-G-HIVLuc infection at multiple steps including, with mechanisms that may differ between M-Mac and GM-Mac. Reduced surface LDLR expression contributed to impaired viral attachment in 27GM-Mac. While additional post-entry restrictions may account for the marked suppression of viral reporter expression. Further studies are needed to establish the contribution of LDLR regulation to this phenotype and to distinguish direct impact from indirect effects of IL-27. Furthermore, given the reported effects of IL-6 and our findings with IL-27 on LDLR expression, it would be of interest to define whether other cytokines that signal through gp130 similarly regulate LDLR expression and susceptibility to viruses that use LDLR for entry and thereby influence susceptibility to VSV-G-pseudotyped viral vectors used for gene delivery.

## Acknowledgement

The authors thank Veronique Nussenblatt for supporting this project. The content of this publication does not necessarily reflect the views or policies of the Department of Health and Human Services, nor does mentioning trade names, commercial products, or organizations imply endorsement by the U.S. Government. This research was supported (in part) by the National Institute of Allergy and Infectious Diseases. The pHIVNL4-3 ΔEnv/Vpr Luciferase were obtained through BEI Resources, NIAID, NIH: Human Immunodeficiency Virus Type 1 (HIV-1) NL4-3 ΔEnv Vpr Luciferase Reporter Vector (pNL4-3.Luc.R-E-), HRP-3418.

